# Model building in SHELXE

**DOI:** 10.1101/2022.04.28.489939

**Authors:** Isabel Usón, George M. Sheldrick

**Affiliations:** Structural Biology, Instituto de Biología Molecular de Barcelona (IBMB-CSIC), Barcelona Science Park, Baldiri Reixach 15, 08028 Barcelona, Spain; ICREA: Institució Catalana de Recerca i Estudis Avançats. Pg. Lluis Companys 23, 08010 Barcelona, Spain; Department of Structural Chemistry, Georg-August Universität Göttingen, Tammannstraße 4, Göttingen, 37077, Germany

**Keywords:** Model building, Phasing, density modification, MRSAD

## Abstract

Density modification is a standard step to provide a route for routine structure solution by any experimental phasing method -with SAD and MAD being the most popular ones- as well as to extend fragments or incomplete models into a full solution. The effect of density modification on the starting maps from either source is illustrated in the case of SHELXE. The different modes in which the program can run are reviewed; these include less well-known uses such as reading external phase values and weights or phase distributions encoded in Hendrickson-Lattman coefficients. Typically in SHELXE, initial phases are calculated from experimental data, from a partial model or map, or from a combination of both sources. The initial phase set is improved and extended by density modification and, if the resolution of the data and the type of structure permits, poly-alanine tracing. The trace now includes an extension into the gamma position or hydrophobic and aromatic side chains if a sequence is provided, which is performed in every tracing cycle. Once a correlation coefficient over 30% between the structure factors calculated from such a trace and the native data indicates that the structure has been solved, in all model building cycles sequence is docked and side chains are fitted if the map supports it. The extensions to the tracing algorithm brought in to provide a complete model are discussed. The improvement in phasing performance is assessed using a set of tests.

**Synopsis:** Side chain tracing now completes model building in SHELXE to enhance density modification. All alternative SHELXE modes, using single or combined sources of starting phase information, are described.

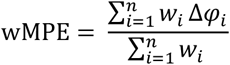

## 1. Introduction

Starting phases from molecular replacement (MR) or experimental phasing (see e.g. Read, 2001 and Hendrickson et al., 1985) are often not accurate enough to make the solution of a macromolecular structure evident and to allow building a complete model for refinement. Still, once a starting solution is obtained it is possible to constrain the electron density to conform to previous structural knowledge. Upon back-transformation of the modified map, combination with the transformed phases rendered is used to improve the original phases. Such procedures are called density modification and were pioneered for macromolecules by Main (Main, 1967) while the first successful application of density modification was reported for small molecules by Hoppe & Gassmann in their phase correction method (Hoppe & Gassman, 1968).

Many sophisticated density-modification schemes have been proposed, and have been incorporated into widely used programs such as DM (Cowtan & Main, 1998), Solomon (Abrahams & Leslie, 1996) and RESOLVE (Terwilliger, 2000). Effective concepts for macromolecular density modification include NCS (non-crystallographic symmetry) averaging (Main, 1967; Bricogne, 1976; Kleywegt & Read, 1997), solvent flattening (Wang, 1985), histogram matching (Zhang & Main, 1990), solvent flipping (Abrahams, 1997) and statistical approaches (Terwilliger, 2000, 2003; Cowtan, 2000). Partial structure interpretation is extremely powerful as it extends information over the whole resolution range and it is widely used, for example in the ARP/ wARP algorithms (Perrakis et al., 2001).

SHELXE implements an alternative approach aiming to enforce stereochemical knowledge, the sphere-of-influence algorithm (Sheldrick, 2002), which is iterated with main-chain tracing (Sheldrick, 2010). In SHELXE, map interpretation has now been extended into the side chains to improve phases and provide a more complete model.

## 2. Density modification in SHELXE

### 2.1. General principles

Figure 1 shows a scheme representing phase improvement by density modification, adapted from the relevant chapter in the International Tables (Zhang, Cowtan & Main, 2001) to illustrate its practical effect in SHELXE. The idea is general, whether starting phases have been determined through experimental phasing -nowadays most frequently SAD or MAD- or calculated from a partial model placed by molecular replacement or through a combination of both sources as in MRSAD (Panjikar et al., 2009). Once approximate phases are available for a structure an electron density map can be computed. This density can be modified based on assumptions on the general physical properties underlying structure: for instance, that in X-ray diffraction density should never be negative (Karle & Hauptman, 1964). Prior knowledge and statistical analysis of its properties are brought into the process. The modified map can be inverted back to calculate structure factors and the resulting phases are expected to have improved. Combining them with the original ones and the experimental amplitudes a new – presumably better– map is calculated to initiate a fresh iteration. Figure 1 illustrates the effect of density modification in the case of a protein originally phased with ARCIMBOLDO (Millán et al., 2015), displaying helices decorating a central beta sheet and containing Zn sites (6YS7). Initial phases are calculated from a single polyalanine helix of 16 amino acids placed by Phaser (McCoy et al., 2007), emphasized in the original map as it provides the only clearly defined feature. As phases are calculated from this helix, the resulting map would in any case show such helical density due to model bias (Brunger, 1990; Luebben & Gruene, 2015), even if the true structure would not contain an helix in this position. The map produced after 20 cycles density modification has developed new features in areas where no initial model was present: in particular, clear density is apparent for a second helix, whereas interpreting the central beta sheet would be more challenging. Eventually, the map after additional model building followed by fresh cycles of density modification becomes very clear, with correct density extending to the side chains. Figure 1b shows the map calculated for the same protein in a different, Zn-containing crystal (6YSD), from the raw SAD phases, after resolving the two-fold ambiguity and adding the heavy atom contribution. In contrast to the initial, model bias dominated, fragment-derived maps in figure 1a, signal is more evenly distributed and noise present everywhere in the map. Figure 1c displays the resulting map after density modification and main chain tracing for this dataset.

**Figure 1.**
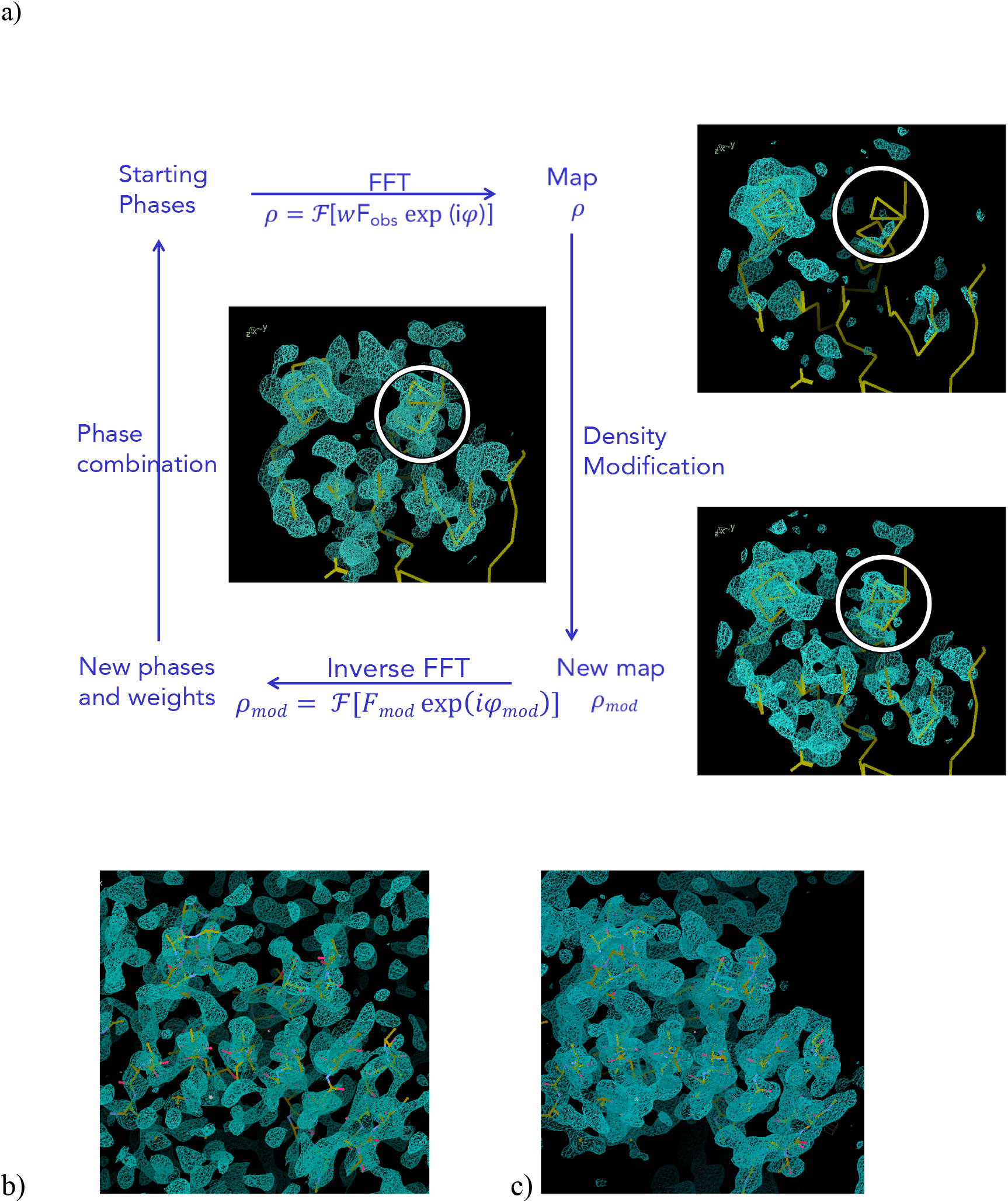
The density modification process (Zhang, Cowtan, Main (2001) International tables of crystallography F, Chapter 15.1. 311) a) exemplified in the extension of a structure from the phases provided by a placed polyalanine helix in the structure of the C-terminal domain of Spr1875 at 1.6 Å resolution. For comparison, maps for the same protein in a different crystal, containing 4 Zn sites (6YSD) are shown. Data collected at a wavelength of 1.28224 Å to a limit of 1.5Å are used. b) Map calculated with raw SAD phases shows everywhere noisy electron density c) The final density-modified map derived from the one shown in panel b.

### 2.2. Initial phase information: the different modes in SHELXE

In practice, SHELXE supports several sources of phase information to be input, be it from an external calculation with a different program or internally generated and combined. Experimental phasing can be exploited for SIRAS, SIR, RIP, MAD or, as in the case summarized in Table 1, SAD, after data preparation with SHELXC. MIR or MIRAS would require an external program, such as Sharp (Vonrhein et al., 2007). Alternatively, phases can be calculated from a (partial) model provided in orthogonal (PDB) or crystal coordinates placed by MR (Thorn & Sheldrick, 2013). Map coefficients, phases and weights in SHELXE format .phi (renamed .phs format), structure factors in .fcf format, generated with SHELXL (Sheldrick, 2015) or phase distributions encoded as Hendrickson-Lattman coefficients (Hendrickson & Lattman, 1970) constitute suitable input. Experimental and map or model phases can be combined, either providing a substructure located e.g. with SHELXD (Usón & Sheldrick, 1999) or ANoDe (Thorn & Sheldrick, 2011) or through an internal cross-Fourier synthesis and peak search. If the substructure is given, SHELXE will refer it to the same origin as the model, and invert it if necessary. It is also possible to perform density modification on the phases derived from model or map and have SHELXE determine the substructure in the final cycle. This last procedure would not combine both sources of phase information but may be useful in the context of some CCP4 pipelines (Winn et al., 2007), for a subsequent iteration or to identify correctly placed partial models.

**Table 1.** SHELXE modes illustrated on a cadmium-containing apoferritin structure.

| Phasing information | Syntax | Parameters | Other files required | MPE ; wMPE: all/centric (°)<br>with main chain tracing |
| --- | --- | --- | --- | --- |
| Experimental (SAD<br>/SIRAS / MAD) | a.phi a_fa | (-h) | a.hkl | 66.8 / 67.3 ; 57.4 / 53.7 |
|  |  |  | a_fa.{res, hkl} | 39.6 / 42.6 ; 28.9 / 23.4 |
| Model pdb | a.pda | - | a.hkl | 72.6 / 73.8 ; 63.8 / 62.7 |
|  |  |  |  | 35.7 / 35.8 ; 26.2 / 19.4 |
| Model crystal<br>coordinates .ins | a | - | a.{ins, hkl} | 72.1 / 72.1 ; 63.6 / 61.6 |
|  |  |  |  | 35.8 / 36.7 ; 26.0 / 19.3 |
| Map coefficients | a.phi | - | a.{ins, hkl} | 72.8 / 72.9 ; 64.6 / 63.2 |
|  |  |  |  | 38.6 / 38.7 ; 27.9 / 20.0 |
| Model + experimental | a.pda a_fa | (-h) | a.hkl |  |
| Given substructure |  | -z0 | a_fa.{res, hkl} | 57.0 / 57.1 ; 45.6 / 39.3 |
|  |  |  |  | 37.5 / 40.0 ; 27.8 / 22.4 |
| Locate and use<br>substructure |  | -z(n) | a_fa.{ins, hkl} | 33.6 / 37.7 ; 24.5 / 20.2 |
|  |  |  |  | 32.3 / 36.1 ; 23.5 / 19.0 |
| Locate substructure |  | - | a_fa.{ins, hkl} | 72.6 / 73.8 ; 63.8 / 62.7 |
|  |  |  |  | 35.7 / 35.8 ; 26.2 / 19.4 |
| Map + experimental | a.phi a_fa | (-h) | a.hkl |  |
| Given substructure |  | -z0 | a_fa.{res, hkl} | 58.3 / 58.5 47.1 / 42.3 |
|  |  |  |  | 37.7 / 40.8 ; 27.9 / 22.7 |
| Locate and use<br>substructure |  | -z(n) | a_fa.{ins, hkl} | 33.2 / 36.9 24.2 / 19.9 |
|  |  |  |  | 32.1 / 35.2 ; 23.4 / 18.6 |
| Locate substructure |  | - | a_fa.{ins, hkl} | 72.3 / 72.6 64.2 / 63.2 |
|  |  |  |  | 36.1 / 36.9 ; 26.3 / 19.3 |
| Hendrickson-Lattman<br>coefficients | a.hlc | - | a.{ins, hkl} | 48.8 / 53.0 ; 39.1 / 38.2 |
|  |  |  |  | 34.9 / 37.4 ; 27.0 / 22.7 |
| Structure factors | a.fcf | - | a.hkl | 72.5 / 72.4 ; 63.7 / 60.6 |
|  |  |  |  | 38.3 / 37.6 ; 27.8 / 20.0 |

Using data from: (a) Mueller-Dieckmann *et al.* (2007). In addition to the mode-specific parameters quoted in the table, all runs have set 20 cycles autotracing and 60% solvent content (`-m20 -s0.6`). A full list and description of parameters is obtained typing *shelxe* in a terminal. For main chain autotracing, three cycles and use of helical seeds were indicated as `-a3 -q`. Fragment optimisation (`-o`) is used for .pda starts. As the data being phased contain Cd, `-h` is given whenever the anomalous substructure is used. All sites and occupancies for the Cd cations were given as in the deposited pdb (2G4H). The tests include a map calculated from this same helical model in SHELXE, without further modification and structure factors in .fcf format, generated with SHELXL (Sheldrick, 2015). Hendrickson-Lattman coefficients were generated in AutoSharp (Vonnrhein *et al.*, 2007) refining the sites and occupancies for the Cd cations in the PDB entry.

The various modes to run the program are all illustrated for the case of an apoferritin structure (PDB id 2G4H) (Mueller-Dieckmann et al., 2007). Results are summarized in Table 1. The single dataset, to a resolution of 2Å contains anomalous signal from the cadmium cations present. An helical polyalanine fragment of 32 residues was placed with Coot (Emsley et al., 2010) as a partial model fragment to provide phases from the derived coordinates and temperature factors, map and structure factors. From the results shown in Table 1, it can be seen that the substructure used in the SAD experiment, reflecting the sites and occupancies in the deposited PDB entry can be improved for phasing purposes. Actually, better results are obtained with fewer atoms and the modes locating the substructure from the partial model or map return the four major Cd sites. SHELXE can also refine the anomalous substructure provided (-z). In addition, it is possible to read in phase distributions as the one shown in the table for this example, which was generated with Sharp starting from the same substructure sites. Anisotropic refinement of the substructure, accounting for various types of scatterers and starting from a phase distribution may be required for difficult cases. The values in Table 1 show how this improved start leads to a much better map upon 20 cycles density modification. In addition, to solve a structure from a MIRAS experiment, this route should be necessary.

A partial model provides an alternative start, with slight differences between pdb and map as the map is used as provided whereas the pdb is trimmed to optimize the correlation to the native data of the structure factors calculated from it. Also default values for sharpening vary slightly for the orthogonal and crystal coordinate formats, due to their typical use contexts.

MRSAD combination of experimental with model or map phases yields the best results. Any of these starts leads to the equivalent solved structure when model building is performed and Table 1 shows the resulting phase errors when three cycles main chain tracing of the map are used to improve phases.

### 2.3. The sphere of influence algorithm

Classical density modification, as established by B. C. Wang (1985) divides the map into protein and solvent regions. Protein regions present the largest density fluctuations and their features should be further enhanced, whereas the density in the more featureless solvent region should be low and uniform and is accordingly flattened. Unique to SHELXE, the *sphere of influence algorithm* (Sheldrick, 2002), avoids locating and smoothing the boundary between protein and solvent. In this algorithm, the variance V of the density is calculated for a spherical surface of radius 2.42 Å (a typical 1,3 distance in a macromolecule) around each voxel in the map. For voxels with a low variance as would be expected within the solvent region, the density at the voxel is ‘flipped’ (ρ’ = -γρ, where γ is typically 1.1 but may be set by the user). The procedure is related to the γ-correction (Abrahams, 1997), except for not requiring an explicit solvent boundary. For voxels with a high variance, typical for protein regions, density is reset to zero if negative and is otherwise left unchanged or subject to a sharpening function (ρ_mod_ =[ρ^4^/(ν^2^σ^2^(ρ)+ρ^2^)]^½^ with ν being resolution dependent, larger the higher the resolution. This function is similar in its behavior to the one used in ACORN (Foadi *et al*., 2000). For intermediate values of the variance, SHELXE applies a weighted mean of the corrected values for the protein and the solvent regions. Sharpening is particularly effective at high resolution or for experimental phases, and in default use it is downweighted for fragment-derived phases.

### 2.4. Extension of partial structures

Mainchain interpretation was incorporated into SHELXE to support phase improvement (Sheldrick, 2010) and extended to include secondary and tertiary structure constraints for lower resolution purposes (Usón & Sheldrick, 2018). SHELXE would typically trace one third to one half of the final structure, to avoid compromising on accuracy because deviations from the correct structure tend to quench the extension process. The factors underlying the chances of success in the extension of a partial structure with given diffraction data are well understood, if not predictable in a quantitative way. They are illustrated in figure 2 through four contrasting examples: The structure of aldose reductase (4LBS) shown in Fig. 2a, in complex with the bromine containing ligand NADP+ and {2-[(4-bromo-2,6-difluorobenzyl)carbamoyl]-5-chlorophenoxy}acetic acid is, at 0.76 Å, one of the highest resolution structures deposited in the PDB (Fanfrlik et al., 2013) for the comparatively large contents of the asymmetric unit: 316 amino acids. Nevertheless, the bromine atom -which can be placed from the native Patterson (Patterson, 1935)-suffices to expand 85% of the structure. Figure 2a illustrates how although initial phases are characterised by an extremely high weighted mean phase error (wMPE) above 80°, even density modification alone succeeds in very slowly improving the phase information, so that after 500 cycles, average phase errors have come down by ∼5°. Main chain autotracing accelerates convergence, and two rounds, interspersed with 10 cycles density modification, bring the errors down to 35°. The subsequent density modification brings the wMPE down to 15° (not shown in the figure). This constitutes a residual difference given that the final deposited structure used as reference contains a model accounting for features established in the course of a high resolution refinement, outside the scope of the model used in phasing: hydrogen atoms, (anisotropic) displacement parameters, multiple conformations for disordered regions, bulk solvent correction and scaling. Phase differences to the deposited structure are therefore never expected to be zero after density modification.

**Figure 2.**
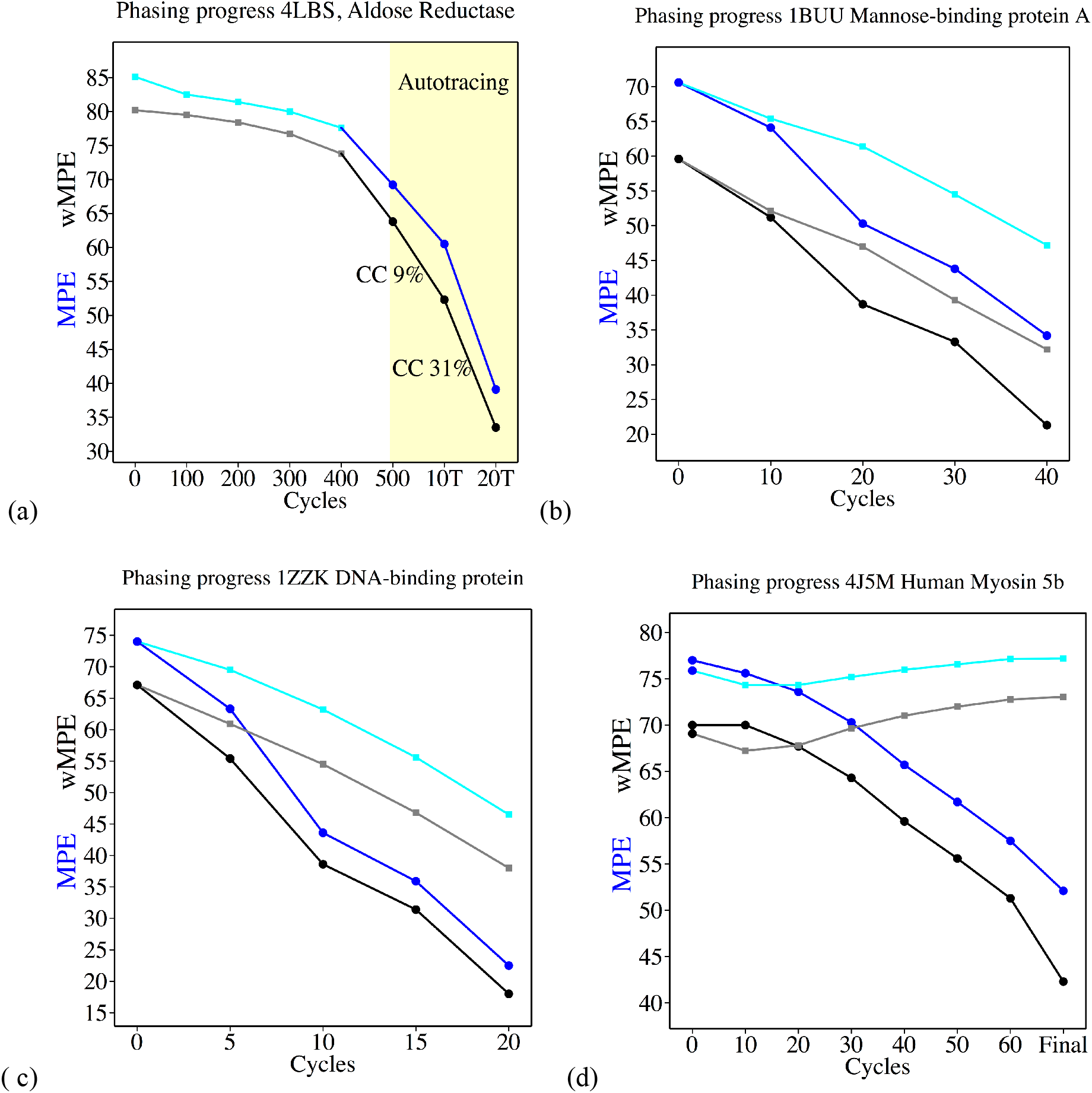
Mean phase error (blue) and weighted mean phase error (grey) starting from fragments and role of model building. Dots show cycles with autotracing, while square points represent no autotracing. a) Aldose Reductase (4LBS, Fanfrlik et al., 2013) at 0.76Å starting from the phases provided by the bromine atom present in the ligand. The deposited structure contains 2567 protein and 687 ligand and water non-hydrogen atoms. After 500 cycles density modification a second run (yellow) iterates autotracing and density modification (10 cycles). b) Mannose-binding protein A of 167 residues at 1.93Å (1BUU, Ng et al., 1998) from a holmium cation. The darker lines with dots (main chain tracing) show a faster convergence and better final phases for the same total number of density modification cycles than when no tracing is applied. c) 1ZZK (Backe et al., 2005) at 0.95 Å, where a 82 residues large structure can be obtained starting from 4 helical amino acids, whereby autotracing is not essential but supports convergence and d) 4J5M human myosin 5b (Nascimento et al., 2013) at 2.07 Å, where main chain interpretation is essential to extend from the two starting helices of 17 alanines to an interpretable map, without autotracing phases deteriorate rather than improve.

The resolution yielded by these aldose reductase crystals is extremely unusual, so in contrast 1BUU exemplifies a structure diffracting to a more typical resolution of 1.93Å, which can also be extended from a single atom to its 150 independent amino acids. Solution starts from a holmium (III) cation to provide starting phases characterised by an already remarkably low wMPE, below 60° (Fig. 2b). It should be remarked that Ho(III), with 64 rather than 36 electrons as in Br^−^, represents a considerable contribution to the total scattering. In Figure 2b, a steeper convergence can be observed during the 40 cycles displayed in the SHELXE run where main chain tracing is used along with density modification versus the run where only density modification is used. This difference becomes negligible if both processes are allowed to run for more cycles, until convergence as in this and many other cases, the final result will be limited by the quality of the data rather than by the starting phase information.

Modern synchrotrons and in-house diffractometers should be able to extract useful anomalous signal for experimental phasing, rendering the first two examples somewhat academic. Nevertheless, in the absence of heavy atoms the same pattern can be seen. Four residues – barely an alpha-helix turn-suffice, when correctly placed, to phase 1ZZK (0.95Å). As seen in figure 2c, in this case tracing accelerates convergence even though in its absence, the same very low overall errors are eventually reached when more cycles are performed (not shown in the figure). Incomplete models at medium resolution may require autotracing for extension into a full solution, 10-15% of the mainchain are typically enough at resolutions around 2Å, as exemplified (Fig. 2d) in the case of human myosin 5B 4J5M (Nascimento *et al*. 2013). This protein is 396 residues long, data extend to 2.1Å and it can be phased from two helices of 17 residues each whereas without building the mainchain model, these starting phases deteriorate and no solution is achieved. Cases like the ones described constitute a frequent target in pipelines such as AMPLE (Bibby *et al*., 2012; Rigden et al., 2018, Simpkin et al., 2019), MRBump (Keegan *et al*., 2018) or ARCIMBOLDO_LITE (Sammito *et al*., 2015).

Availability of high resolution data has so far been critical for the extension of features outside the placed partial structures when these constitute a very limited fraction of the contents of the asymmetric unit. Some improvement is achieved extrapolating unmeasured data whether missing low resolution data or reflections beyond the resolution limit (Usón *et al*., 2007), in what has been named the “free lunch” algorithm. This option is used for generating electron density maps and can produce spectacular results for high resolution and/or high solvent content.

### 2.5. No structure solution despite partially correct phase information

As suggested in this last example, borderline cases occur, where despite partially correct start phases it may still not be possible to solve the structure interpreting the experimental map, extending a partial structure or eliminating the errors in a correctly placed model presenting large geometrical differences to the target. Locating the anomalous substructure or solving the molecular replacement problem is not necessarily equivalent to solving the structure. When the starting phases are not accurate enough –the more so the poorer the resolution– the correct starting information cannot be successfully extended and non-random starts remain unsolved. Also, a high percentage of helical structure is advantageous versus predominantly beta structures. In such cases, the brute force method implemented in SLIDER of probing all favourable side chain assignments onto a trace can extract a solution (Borges *et al*., 2020). Our experience with this program underlies the choices to extend the model towards the side chains in the SHELXE implementation. Other approaches such as the sophisticated combination of building and refinement developed over the last three decades in ARP/wARP (Chojnowski *et al*., 2020) should also be mentioned here. In the case of SLIDER, we observed that in the absence of powerful hardware to support the heavy calculations associated to probing all possible side chain assignments, simplified modes considering only aromatic and hydrophobic residues or even reducing every side chain to a serine (Schwarzenbacher *et al*., 2004) occasionally allowed to pull a complete solution from a poor start.

### 2.6. Gamma extension and map probing

Even in the absence of sequence information it can be safely assumed that most residues will have a sidechain with a carbon, oxygen or sulphur atom in gamma position. As the density modification proceeds, main chain electron density tends to be revealed earlier and more prominently but even with large mean phase errors (see Figure 1) clear electron density starts showing in parts of the structure. Gamma positions typically cluster in one of three staggered positions (Lovell *et al*., 2000). Probing density in each of these at a compromise distance of 1.47Å, between the shorter CB-OG in serine and the CB-CG distance in most amino acids establishes if the difference between the highest and lowest electron density value is significant. Furthermore, it allows the detection of features in the map. Every auto tracing cycle, the trace is probed at the gamma position, provided there is clear density for the beta position. Otherwise, the residue is annotated as probable glycine. If there is clear discrimination between the highest and lowest density in the alternative position, the residue is modelled as pseudo-serine, with a slightly longer distance. Fig. 3a illustrates the inclusion of some gamma positions in the trace of a map for 4ICI, when it still has large MPE above 70°. If the maximum is not in trans, the +/-30° conformation particular to proline will be probed with the appropriate geometry for its gamma carbon (Fig. 3b). Finally, if the intermediate and highest density values are similar and clearly discriminated from the lowest one, the residue is annotated as probable valine, threonine or isoleucine. Gamma sites are included in the calculation of trace-derived phases for the next cycle. The improvement this brings is modest in the first cycle, less than 2° in the best cases, but in no case has it been found to deteriorate the phases, as Table 2 shows for a set of test cases.

**Figure 3.**
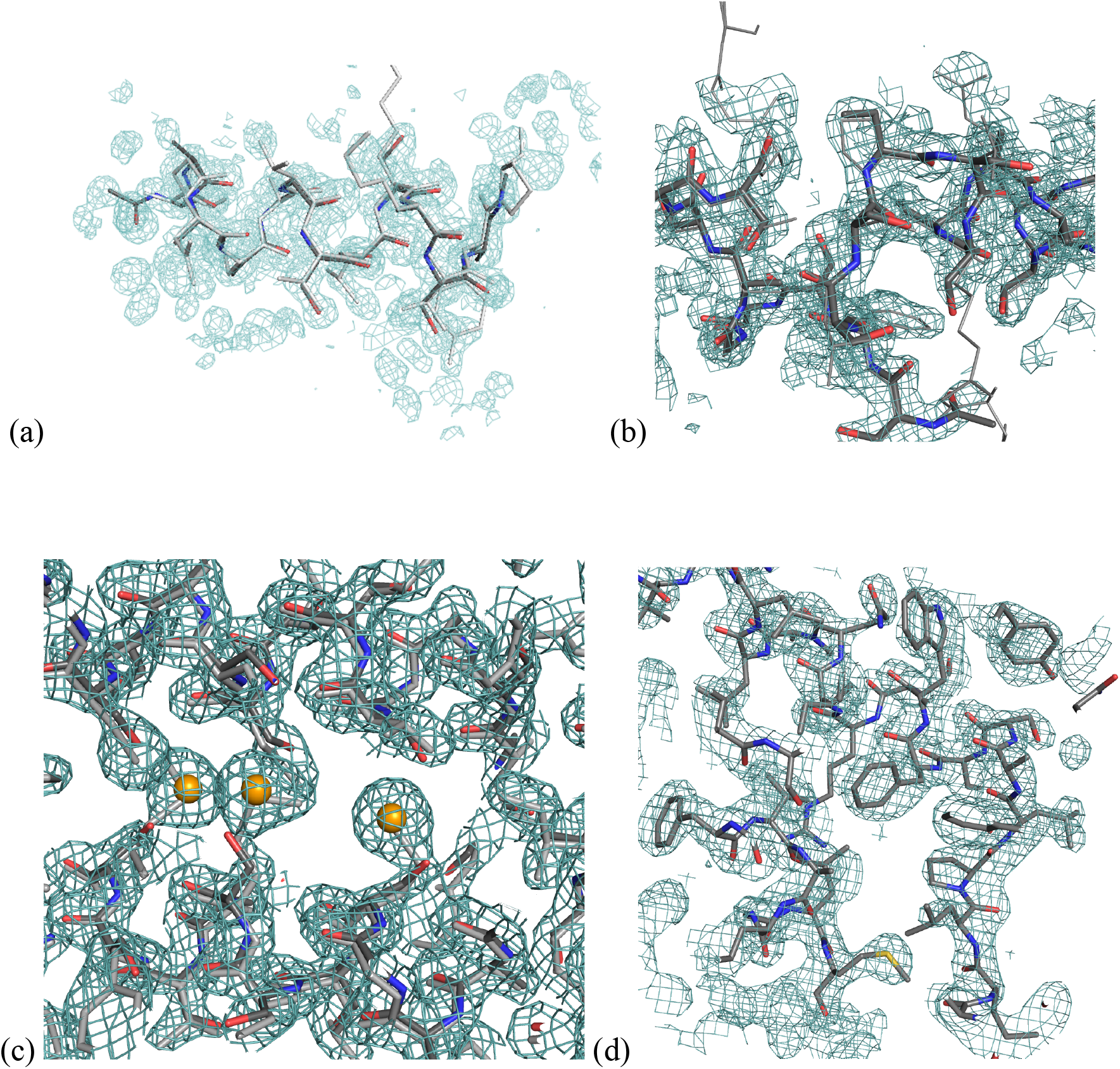
Tracing into the side chain density. a) Tracing of gamma positions illustrated in the first cycle of 4ICI, starting from experimental phases provided by two selenium sites. b) proline is identified from the density and position in the trace. c) Use of substructure sites as sequence markers. d) Traced model with sidechains for CAS3.

**Table 2.** Summary of the effect of gamma tracing on test structures.

| ID | Start | Space<br>Group | d(Å) | CC<br>– / –O | Parameters | MPE (wMPE)<br>– / –O |
| --- | --- | --- | --- | --- | --- | --- |
| hirustasin | 10S | P4<sub>3</sub>2<sub>1</sub>2 | 1.2 | 46.8 | –m25 –s0.4 –a1 –K | 41.9 (35.1) |
|  | fragment |  |  | 47.5 |  | 41.3 (34.7) a |
| insulin | 6S SAD | I2<sub>1</sub>3 | 1.54 | 37.4 | –m10 –s0.4 –h –a1 | 53.8 (47.7) ; |
|  |  |  |  | 33.3 |  | 51.4 (46.0) b |
| isomerase | 2Mn SAD | I222 | 1.56 | 32.9 | –m10 –s0.49 –h –a1 | 56.7 (49.7) |
|  |  |  |  | 34.8 | –i | 55.4 (48.5) b |
| Spr1875Ct | H14 | P3 | 1.6 | 11.6 | –m100 –s0.5 –a1 – | 71.4 (63.9) |
|  | fragment |  |  | 12.2 | B1 | 71.3 (63.8) c |
| 3L22 | 12Se SAD | P4<sub>3</sub>22 | 1.9 | 39.5 | –m15 –s0.45 –a1 –q | 48.8 (43.2) ) |
|  |  |  |  | 42.3 | –h | 46.7 (40.0) d |
| CAS3 | 2H16 | P4<sub>3</sub>32 | 1.9 | 19.1 | –m15 –s0.5 –d1.9 – | 65.1 (57.7) |
|  | fragment |  |  | 19.6 | a1 –q | 64.8 (57.5)e |
| bucandin | 10 S | C2 | 0.96 | 30.5 | –m25 –s0.4 –a1 –K | 56.5 (48.7) |
|  | fragment |  |  | 30.7 |  | 55.9 (48.2)f |
| Proteinase<br>K | map | P4<sub>3</sub>2<sub>1</sub>2 | 1.27 | 12.8 | –m15 –s0.5 –a1 – | 70.3 (64.0) |
|  |  |  |  | 13.4 | S3 –e1 –I15 | 69.9 (63.4)g |
| amiA | Polyalanine | P2<sub>1</sub>2<sub>1</sub>2<sub>1</sub> | 1.2 | 35.0 | –m50 –s0.5 –a1 –q | 45.0 (37.6) |
|  | fragment |  |  | 36.8 | –e1 –I50 –v0.1 | 43.5 (36.1)h |
| 4ICI | 3Se SAD | I4<sub>1</sub> | 1.4 | 2.6 | 4ici 4ici_fa –m30 –h | 82.0 (77.6) |
|  |  |  |  | 2.9 | –s0.4 –i –a1 | 82.0 (77.4) |
| PilA1 | 2H30 | P4<sub>1</sub>2<sub>1</sub>2 | 1.65 | 10.6 | –m25 –s0.5 –a1 –q | 76.4 (70.4) |
|  |  |  |  | 10.9 |  | 76.3 (70.4)i |

Using data from: (a) Usón *et al.* (1999), (b) Sevvana *et al.* (2020), (c) Acebrón *et al.* (2020), (d) Vollmar *et al.* (2020), (e) Freitag-Pohl *et al.* (2019), (f) Kuhn *et al.* (2000), (g) Wang *et al.* (2006), (h) Hermoso *et al.* (2022), (i) Crawshaw *et al.*, (2020). CC for the trace, wMPE is the weighted mean phase error for this trace. Parameters used in the SHELXE line: –m number of density modification cycles; –h number of heavy atom sites to use; –s solvent fraction; –a auto-tracing; –q length of helix template, default 7; –K retain input fragment.

### 2.7. Sequence docking

Sequence docking is performed combining experimental evidence from the features in the current electron density map and previous structural knowledge.

Sequence information is input through a fasta format file named with the root name of the data file and the extension .*seq*. If the flag -O is set but no sequence file is provided the program will issue a warning and perform only gamma tracing to aid phasing, so that pipelines would not fail for lack of this file. To assign the corresponding sequence to each of the traced chains, these are considered from longest to shortest. Then probabilities are calculated for all possible sequence assignment obtained sliding a copy of the sequence, with two trailing dummy residues on each end. Probabilities are calculated as the sum of individual probabilities of each residue in the trace on a logarithmic scale, combining the score obtained probing the electron density and the probability derived from prior structural knowledge, following the scheme described in (McCoy, 2004).

If there is a substructure of anomalous scatterers comprising selenium from selenomethionine or sulfur from cysteine and/or methionine, its atoms are used as sequence markers (Fig. 3c).

There is a vast amount of prior knowledge on sequence-structure relationship but the reliability of secondary structure prediction from the sequence is limited in the absence of comprehensive PDB data (Berman *et al*., 2000, Lange *et al*., 2020). It would be possible to involve the powerful methods implemented in HHPRED (Söding *et al*., 2005) as an external dependency but rather than a comprehensive sequence analysis, SHELXE sequence assignment is guided by the more robust principles, which stuck out clearly enough to be identified when still a very scarce subset of protein structures had been determined. Notably, the overall propensities of some amino acids to secondary structure (Chou and Fassman, 1974), particularly the residues typically terminating /initiating secondary structure elements or marking loops (Richardson & Richardson, 1988) (e.g. the proline displayed in Fig. 3b) and the consistent association of hydrophobic residues upon sequence docking to the trace of a strand or an helix (Eisenberg *et al*., 1983). Fundamentally, it is the available electron density map that can be interrogated and the secondary structure of the trace internally described with characteristic vectors (Medina *et al*., 2020).

### 2.8. Error correction

As sequence docking relies on correct main chain tracing, a previous step to locate and remove connections of unusual Ramachandran values (Hollingsworth & Karplus, 2010) and lower density than that of flanking connections has been introduced to precede sequence docking. In general, to avoid errors the criterium used to accept an assignment is that it needs to be distinctly better than any alternative. Before incorporating side chains into the final trace (Fig. 3d), a comparison of the CC calculated omitting the side chains for every stretch of chain is performed, analogous to the PDB optimization step introduced in SHELX macromolecular phasing (Sheldrick & Gould, 1995). If the CC characterizing the polyalanine trace is higher than when sidechains are incorporated, they are eliminated from that part of the model.

## 3. Tracing tests

Figure 4 displays the results of the phasing and model building of apoferritin (Müller-Dickman et al., 2007) previously described (2.2) starting from anomalous difference data and Cd sites or phase probability distributions derived thereof as Hendrickson-Lattman coefficients or alternatively from an equivalent fragment of a long helix provided as a pdb file or in fractional crystal coordinates, as well as structure factors or a map calculated from the same model. Combinations of SAD and fragment or map phases are also included. The last model building cycle involves side chain tracing. In all these cases, results after three groups of 20 cycles density modification interspersed with map tracing are comparable. Nevertheless, in the cases where more complete starting information -combining model/map and SAD phases- is used convergence is faster and a more complete trace is present already in the first cycles.

**Figure 4.**
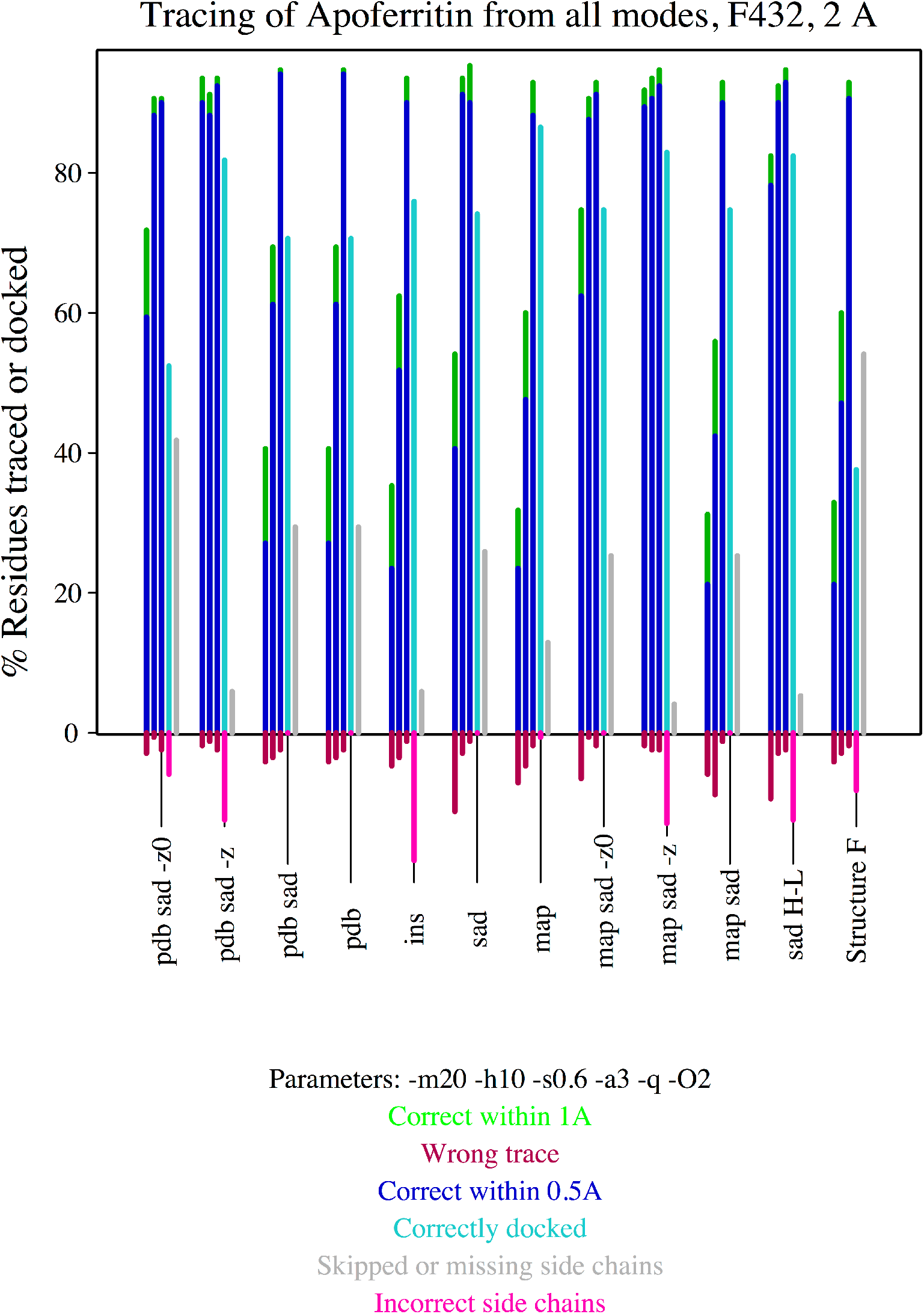
Percentage of main-chain correctly traced within 0.5Å (blue), within 1Å (green), incorrect trace (red) during the three autotracing cycles and performance on the side chain tracing in the final cycle as percentage docked (cyan), skipped or absent from the main chain trace (gray) and incorrectly traced (pink) for the apoferritin (2G4H) starting from experimental, model or combined phases.

Furthermore, eleven structures with resolutions ranging from 1.2 to 2.0 Å. have been used to test and illustrate the new side chain tracing features in SHELXE described above. Table 3 summarizes the characteristics, parameterisation and phasing results obtained incorporating side chain tracing in SHELXE for SAD phasing and fragment cases where the atoms in the substructure or starting structure can be used as marker as well as when this is not the case. For molecular replacement solutions, initial phases should be limited (with the parameter -y) to a resolution dictated by the rmds and extended in the course of density modification, the more limited the lower the identity between template and structure. For fragment phases, this default should be changed to use the full resolution available as small fragments should be accurate to be able to solve a structure. Hirustasin (Usón *et al*., 1997) and bucandin (Kuhn *et al*., 2000) have each been extended from 10 sulfurs; SAD data for insulin and glucose isomerase were measured from non-merohedrally twinned crystals (Sevvana *et al*., 2019). The SusD protein 3L22 was originally solved through a MAD experiment (Vollmar *et al*., 2020); for the purpose of this study it was phased from the peak wavelength used as SAD data at a resolution of 1.9Å. VirusX CAS3 (Freitag-Pohl *et al*., 2019) was the first previously unknown structure where we used the SHELXE development version to build the model with side chains. The flavoprotein 4ICI in I4_1_ was incorporated as a test where space group inversion has to be performed at (0.5, 0.25, 0.5) rather than at the origin, an operation performed when phases from the fragment and from the pre-calculated anomalous substructure need to be referred to the same origin. Proteinase K (Wang *et al*., 2006) and amia (Hermoso *et al*., 2022) are proteins phased from fragments of homologs, placed and refined with ARCIMBOLDO_SHREDDER (Millán *et al*., 2018). The first is a test case starting from a map as ALIXE (Millán *et al*., 2020) combines solutions in reciprocal space and outputs a set of phases and figures of merit. The second one was originally solved with SHREDDER and the test starts from a fragment.

**Table 3.**
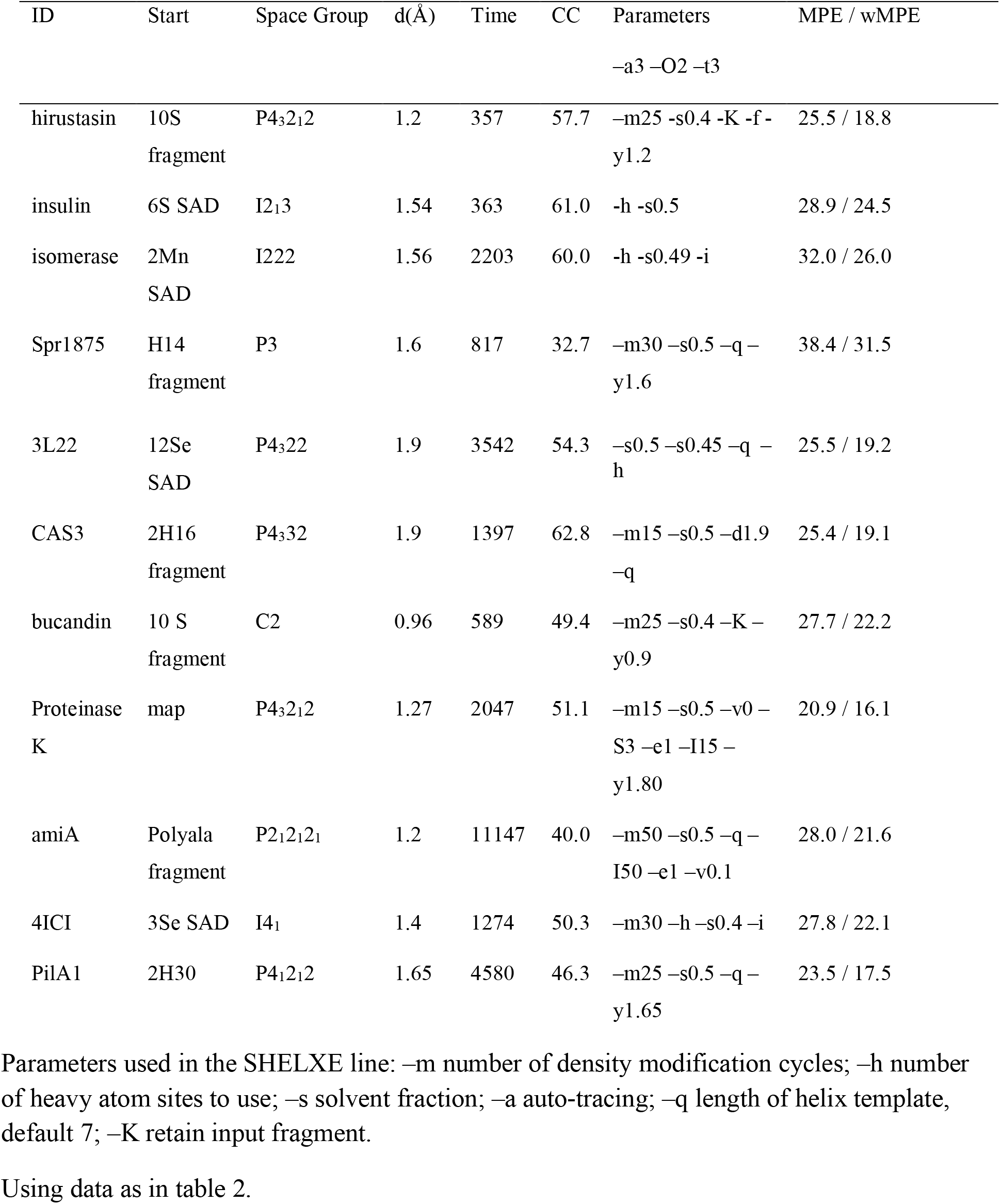
Summary of density modification and model building with SHELXE on test structures.

Finally, PilA1 (Crawshaw *et al*., 2020) was originally solved with ARCIMBOLDO_LITE and contains three chains of 150 amino acids. For structures like this one, with NCS, the fasta format file read by SHELXE should explicitly contain a copy of the sequence for each of the chains present. The tracing results for all these cases are shown in Figure 5.

**Figure 5.**
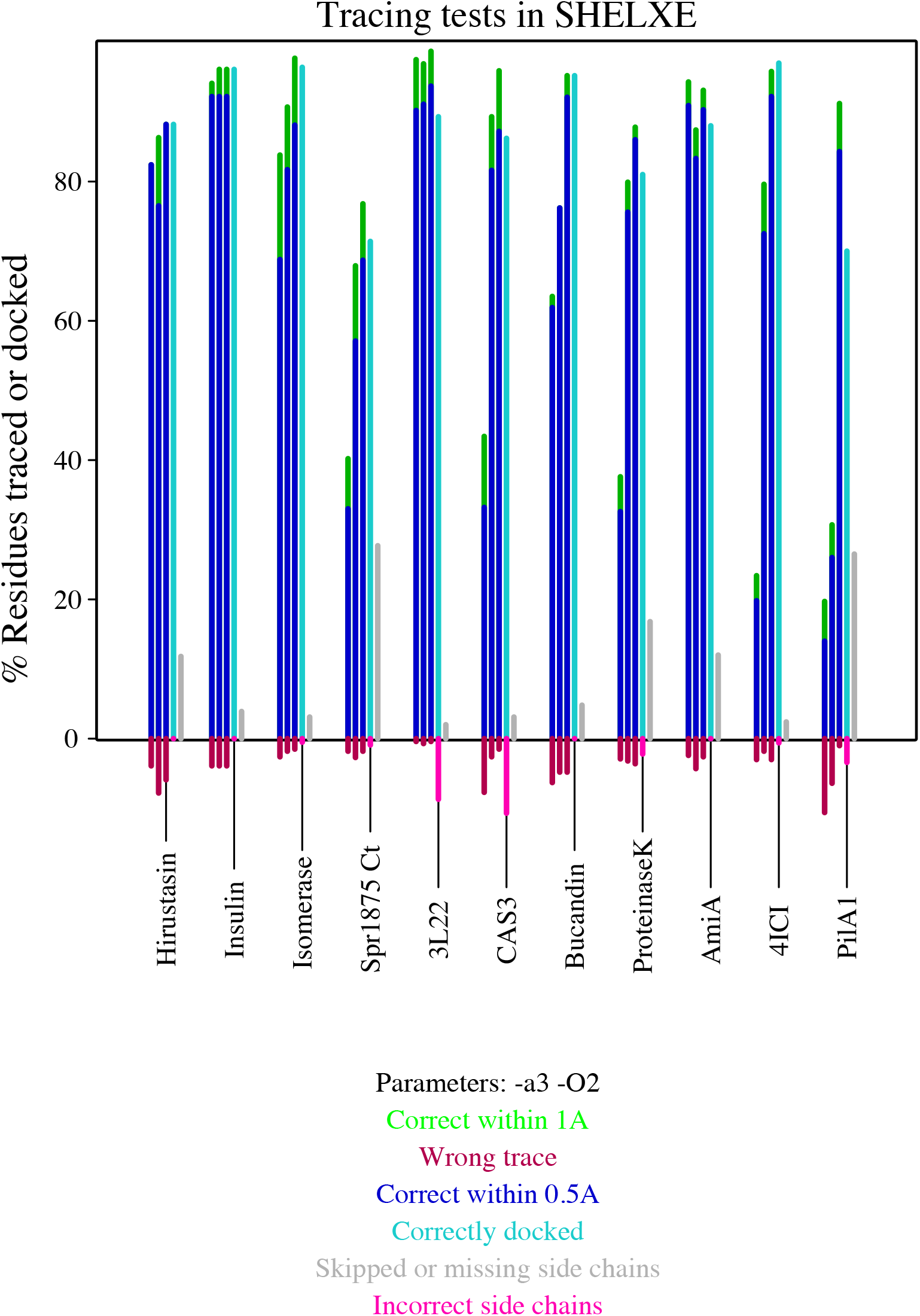
Percentage of main-chain correctly traced within 0.5Å (blue), within 1Å (green), incorrect trace (red) during the three autotracing cycles and performance on the side chain tracing in the final cycle as percentage docked (cyan), skipped or absent from the main chain trace (grey) and incorrectly traced (pink) for the structures summarised in tables 2 and 3.

## 4. Concluding remarks

This paper provides an overview of model building in SHELXE and of its effect as a constraint on density modification. It also describes all the different modes in which SHELXE can be used and showcases how density modification can have a decisive contribution on phasing, which is sometimes hidden within pipelines.

The tracing algorithms have been expanded to enhance performance in borderline cases and to provide more complete models. Thus, incorporation of side chain atoms extends the phasing improvement brought about by model building. Furthermore, obtaining a more complete model, with side chains assigned and fit to the density is convenient. In the absence of a sequence or at lower resolution, tracing of poly-serine has been added to SHELXE in all tracing cycles to increase model scattering. If a sequence is provided SHELXE can assign and fit sidechains to the trace, which in the tests presented has been performed in the last tracing cycle, after extension of the gamma position in all previous cycles.

In view of the results presented, the recommended use, corresponding to the default triggered by the flag –O when a sequence is provided is that every autotracing cycle gamma extension and density probing will be performed, incorporating probable side chains for aromatic and hydrophobic residues with partial occupancy of 0.6 to the trace used to generate phases for the next round of density modification. Still, models are output as polyalanine at this stage. Once the CC characterising the trace reaches 30%, sequence docking will be performed in all remaining autotracing cycles, the best scored model with side chains saved as a pdb and its derived phases, combined in the calculation of the output map. SHELXE is often encountered within phasing pipelines, where a correlation coefficient greater than 25% between the structure factors calculated from the poly alanine trace and the native data is adopted as an indication that the structure has been solved for a resolution of 2.5Å or better. As seen in table 3, CC values up to twice those typically rendered by main chain tracing are obtained from the complete models. Therefore, the procedure implemented ensures that the CC value will be consulted on the polyalanine trace and that the most complete and correct model will be output incorporating side chains for a solved structure.

Performance of this procedure was assessed within the auto-rickshaw pipeline on a set of 40 structures not previously used to develop the algorithms. The resolution in this pool of structures ranged between 2.0 and 2.4 Å, yielding improved results over the previous distributed version and nearly complete models.

## Abbreviations

CC: Correlation coefficient expressed as percentage

## Acknowledgements

We thank Clemens Vonrhein for his help generating apoferritin HL-encoded starting phase distribution with Sharp and Santosh Panjikar for thoroughly testing the new model building on an independent pool of structures within his Auto-Rickshaw pipeline and for useful discussion and feedback. We are grateful to the State of Niedersachsen GMS. IU acknowledges grant PGC2018-101370-B-100 (MICINN/AEI/FEDER/UE) and support from STFC-UK/CCP4: “Agreement for the integration of methods into the CCP4 software distribution” and for a Visiting Fellowship at Clare Hall College, Cambridge.

